# AhR ligands from LGG metabolites promote piglet intestinal ILC3 activation and IL-22 secretion to inhibit PEDV infection

**DOI:** 10.1101/2023.12.05.570065

**Authors:** Junhong Wang, Yibo Zhao, Tong Cui, Hongyu Bao, Ming Gao, Mingyang Cheng, Yu Sun, Yiyuan Lu, Jiayao Guan, Di Zhang, Yanlong Jiang, Haibin Huang, Chunwei Shi, Jianzhong Wang, Nan Wang, Jingtao Hu, Wentao Yang, Guilian Yang, Yan Zeng, Chunfeng Wang, Xin Cao

## Abstract

In maintaining organismal homeostasis, gut immunity plays a crucial role. The coordination between the microbiota and the immune system through bidirectional interactions regulates the impact of microorganisms on the host. Our research focused on understanding the relationship between substantial changes in jejunal intestinal flora and metabolites and intestinal immunity during porcine epidemic diarrhea virus (PEDV) infection in piglets. We discovered that *Lactobacillus rhamnosus GG* (*LGG*) could effectively prevent PEDV infection in piglets. Further investigation revealed that *LGG* metabolites interact with type 3 innate lymphoid cells (ILC3) in the jejunum of piglets through the aryl hydrocarbon receptor (AhR). This interaction promotes the activation of ILC3 cells and the production of interleukin-22 (IL-22). Subsequently, IL-22 facilitates the proliferation of IPEC-J2 cells and activates the STAT3 signaling pathway, thereby preventing PEDV infection. Moreover, the AhR receptor exerts its influence on various cell types within organoids, including intestinal stem cells (ISCs), Paneth cells, and enterocytes, fostering their growth and development, suggesting a broad impact of AhR on intestinal health. In conclusion, our study demonstrates the ability of *LGG* to modulate intestinal immunity and effectively prevent PEDV infection in piglets. These findings highlight the potential application of *LGG* as a preventive measure against viral infections in livestock.

## Introduction

Porcine epidemic diarrhea virus (PEDV), a member of the genus Coronavirus A of the family Coronaviridae, causes acute diarrhea and/or vomiting, dehydration, and high mortality in neonatal piglets (Jung and Saif 2015). Related studies have reported significant changes in the intestinal microbiota of *Lactobacillus*, *Escherichia coli*, and *Lactococcus* in piglets as a result of PEDV infection (Yang et al., 2020a). Specifically, the abundance of *Lactobacillus* jejuni and *Lactobacillus* cecum showed a decreasing trend in PEDV-infected piglets, accompanied by significant changes in metabolites(Li et al., 2022).

In recent years, it has been found that intestinal immunity has an important role in organismal homeostasis (Yu et al., 2022). Bidirectional interactions between the microbiota and the immune system coordinate many microbial effects on the host (Ronan et al., 2021). Innate lymphoid cells (ILCs), which are continuously interacting with commensal flora in the gut, are the pioneers (Gury-BenAri et al., 2016). Meanwhile, IL-22 production by ILC is critical for intestinal immunity during the early course of infection development (Satoh-Takayama et al., 2008). Previous studies have shown that intestinal flora is a key factor in IL-22 production in the gut, but the regulatory mechanisms are unclear (Yang et al., 2020b).

In addition to interacting with a variety of intestinal epithelial cells, ILCs also interact with intestinal stem cells (ISCs) in the crypts to regulate the differentiation and function of ISCs (Lindemans et al., 2015). ILC3-derived IL-22 can promote the phosphorylation of signal transducer and activator of transcription 3 (STAT3) on intestinal stem cells through the action of IL-22R, thus inhibiting the apoptosis of intestinal stem cells. At the same time, IL-22 can stimulate the proliferation of intestinal stem cells and then ameliorate the damage to ISCs caused by chemotherapy (Hanash et al., 2012; Aparicio-Domingo et al., 2015).

Aryl hydrocarbon receptor (AhR) is essential for the maintenance of ILC3 and the production of IL-22 (Pernomian et al., 2020). AhR is another key transcription factor for ILC3, and the presence of lymphoid tissue in the intestine is associated with intact AhR expression (Xia et al., 2017). In contrast, the removal of AhR is equivalent to the absence of ILC3, resulting in the disappearance of intestinal lymphoid tissue (Kiss et al., 2011). AhR is expressed in many mammalian tissues, particularly in the liver, intestine, and kidney. In the intestine, AhR, expressed mainly by epithelial cells and innate immune cells, plays an important role in the regulation of innate immunity and regulates the number of lymphocytes within the epithelium. When the energy source is converted from sugar to tryptophan, *Lactobacillus* lactis amplifies and produces indole-3-aldehyde (I3A), which serves as an AhR ligand that promotes the production of IL-22 by ILC3 (Zelante et al., 2013). IL-22 plays a central role in the protection of host cells against pathogens at the surface of the mucosa through the activation of STAT3 (Backert et al., 2014). It has been reported that mature IL-22 of porcine origin inhibits PEDV infection and promotes the expression of antimicrobial peptide β-defensins, cytokine IL-18, and IFN-λ through activation of the STAT3 signaling pathway (Xue et al., 2017).

An important way in which probiotics combat diarrheal diseases is by modulating the mucosal immune system of the host intestine. Mucosal immunity is one of the important components of the body’s immune system, and its main function is to remove pathogenic microorganisms that invade the body through mucosal surfaces. The mucosal immune system has a unique organizational structure and function. It is widely distributed in the mucosal tissues of the respiratory, digestive, and genitourinary tracts, serving as the main site of local immune response. The mucosal immune system consists of various components, including mucosal epithelial tissue, mucosal-associated lymphoid tissue (MALT), intestinal epithelial cells, immune cells, and the molecules or secretions they produce.

Additionally, it includes mucosal microorganisms, such as commensal microorganisms, which normally inhabit the mucosa. Some studies have reported that oral administration of *Lactobacillus* inhibits PEDV infection (Yang et al., 2023). Colonization of mice with the model probiotic *Lactobacillus rhamnosus GG* (*LGG*) enhances the maturation of intestinal function and the production of IgA and produces promising results in protection against intestinal injury and inflammation (Yan et al., 2017).

In this study, a co-culture system of porcine jejunal gut tract-like organoids with porcine jejunal lamina propria lymphocytes (LPLs) was established to examine and validate the protective effect of *Lactobacilli* on intestinal epithelial cells. Our hypothesis suggests that *LGG* may secrete substances, such as I3A, which can activate the AhR receptors located on the surface of ILC3 present in the lamina propria of the pig intestinal mucosa. This activation could potentially enhance the secretion of IL-22 by ILC3. Then, the STAT3 signaling pathway of intestinal epithelial cells was activated, regeneration of ISCs was induced to perform damage repair function, Paneth cells were induced to secrete Reg3b and Reg3g to inhibit the growth of pathogenic bacteria, and intestinal cells were induced to secrete IFN-λ to inhibit PEDV replication.

## Materials and Methods

### Animals and sample collection

We used 3-day-old SPF piglets, which were purchased from the Harbin Veterinary Research Institute. All piglets were transported in enclosed carts designed for SPF conditions. We collected samples from the jejunum and other tissues of the piglets. Then, we analyzed the immune cells in the jejunal tissues using flow cytometry, processed and analyzed the samples using q-PCR and other experimental methods, and sectioned and stained the tissue samples. All animal experiments complied with the requirements of the Animal Management and Ethics Committee of Jilin Agricultural University.

### Cell separation

Cell samples obtained from the jejunum of piglets were subjected to subsequent flow assay and in vitro cell culture and q-PCR. Firstly, after euthanasia of piglets, about 5 cm of jejunum was dissected longitudinally, rinsed with PBS and divided into 1 cm sized intestinal fragments, which were then transferred to the separation solution (15 mL of RPMI-1640, 1% penicillin and streptomycin, 1% HEPES, 5 mM EDTA, 2 mM DTT, and 2% heat-inactivated fetal bovine serum (FBS)) and incubated for 28 minutes in a shaking incubator at 37°C and 200 rpm, and then removed. After incubation for 28 minutes in a shaking incubator at 37°C and 250 rpm, the intestinal fragments were obtained rinsed and continued into the enzyme digestion solution (8 mL RPMI-1640 medium, 1% penicillin and streptomycin, 1% HEPES, 50 mg collagenase IV, 1 mg DNase I, and 2% FBS), and incubated for 28 minutes in a shaking incubator at 37°C and 250 rpm before being removed, and then filtered through a 70-μm cell strainer to get the LPL cells in the jejunum of piglets.

### Antibody Information

Invitrogen: CD2 Monoclonal Antibody (14-0029-82), CD3e Monoclonal Antibody (MA5-28774), CD3e Monoclonal Antibody (Biotin) (MA5-28771), CD11b Monoclonal Antibody (MA5-16604), CD11c Monoclonal Antibody (APC) (17-0116-42), CD21 Monoclonal Antibody (A1-19243), CD21 Monoclonal Antibody (PE) (MA1-19754), CD45 Monoclonal Antibody (FITC) (MA5-28383), CD117 (c-Kit) Monoclonal Antibody (APC) (17-1171-82), CD163 Monoclonal Antibody (PE) ( MA5-16476), CD172a Monoclonal Antibody (MA5-28299), CD335 (NKp46/NCR1) Monoclonal Antibody (PE) (MA5-28352), SLA Class II DR Monoclonal Antibody ( MA5-28503), Gata-3 Monoclonal Antibody (PE-Cyanine7) (25-9966-42), T-bet Monoclonal Antibody (PE) (12-5825-82), Goat anti-Mouse IgG H&L (PE-Cyanine5) (M35018).

BD Pharmingen: Fixable Viability Stain 780 (565388), Purified Rat Anti-Mouse CD16/CD32 (Mouse BD Fc Block) (553142), Rat Anti-Pig γδ T Lymphocytes (PE) (551543), Rat Anti-Pig γδ T Lymphocytes (561486), Mouse anti-GATA3 (558686).

Abcam: Streptavidin protein (Alexa Fluor 594) (ab272189), Streptavidin protein (APC) (ab243099), Goat Anti-Mouse IgG H&L (FITC) (ab6785), Goat Anti-Rabbit IgG H&L (Alexa Fluor 488) (ab150077), Goat Anti-Mouse IgG H&L (Alexa Fluor 488) (ab150113), Goat Anti-Rat IgG H&L (Alexa Fluor 488) (ab150165), Goat F(ab’)2 Anti-Mouse IgG H&L (PE) (ab7002), Goat F(ab’)2 Anti-Rat IgG H&L (PE-Cyanine5) (ab130803), Goat Anti-Mouse IgG H &L (Alexa Fluor 647) (ab150115), Rabbit Anti-Goat IgG H&L (Alexa Fluor 488) (ab150141), Rat Anti-Mouse IgG1 H&L (PE) (ab99605). Anti-Villin antibody (ab244292). Anti-EpCAM antibody(ab71916). Anti-EpCAM antibody(ab71916). Anti-Ki-67 antibody(ab15580). Anti-LGR5 antibody(ab273092).

Solarbio: CFSE (S1076), PMA (P6741), D-PBS (D1040), PBS (P1010)
GE Healthcare: Percoll (17089101).
R&D: APC Anti pig IL-4 (1644057).

Sigma: Penicillin And Streptomycin (V900929), Collagenase IV (V900893-1G), Ionomycin (56092-81-0), DNase I (10104159001), DTT (3483-12-3), HEPES ( H3375), EDTA (E8008), FBS (F8318).

PE Goat anti Mouse(SPP101), FITC Rabbit anti Goat (SPP101-100), PE anti Mouse (SHP501) purchased from 4A Biotech.,Ltd.

### Preparation of IL-22 protein and IL-22 monoclonal antibody

The porcine IL-22 (Gene ID: 595104) gene was synthesized by consulting NCBI, and the structural domains and were examined to determine the protein expression scheme, and the prokaryotic and eukaryotic expression vectors were prepared, and the IL-22 protein was obtained after the prokaryotic expression was purified, and the IL-22 protein was immunized to the mice and then taken from the B-cells for the fusion of hybridoma cells, and large quantities of monoclonal antibodies were prepared by means of ascites preparation. A large number of monoclonal antibodies were prepared by ascites preparation, and the antibodies were labeled with antibody labeling kit (ab201795) after purification. This part of the work was done with the help of Shanghai Company.

### LGG and EVs Acquisition

LGG (ATCC 53103) was grown in De Man, Rogosa, and Sharpe (MRS) broth for 12 h at 37°C. After culturing overnight, the bacteria were inoculated 1:100 in fresh MRS broth and grown under anaerobic conditions until reaching the mid-log phase. Then, the colonies were counted, and the cell density was adjusted to 5 × 10^9^ colony-forming units (CFU)/mL.

The culture was centrifuged overnight at 7,000 g for 30 min to eliminate debris, including dead cells and other waste products. The supernatant obtained was filtered and ultracentrifuged at 150,000g for 70 min. The precipitate was collected after centrifugation (LGG-EVs were mainly enriched in the precipitate, with minimal presence in the supernatant) and stored at −80°C for later use.

### The PEDV information

The PEDV strain (PEDV LJX01/2014, PEDV N gene GenBank number: MK252703) was provided by the Professor Guangliang Liu, Lanzhou Veterinary Research Institute, China. The propagation and titration of the PEDV LJX strain were described previously (Chen et al., 2020). Piglets were infected by oral feeding using a virus with a copy number of 1.35 × 10^4^ RNA copies/g, and each piglet was infected with 5 mL of virus solution.

### Experimental animal models

Three-day-old piglets were treated with oral antibiotics, in which gentamicin 3 mg, metronidazole 20 mg, and ampicillin 100 mg per kg of piglets were fed for 7 d. The CON group was fed for PBS 14 d as a control group, the ABX + LGG + PEDV group received PEDV infection after 7 d of antibiotic feeding and further LGG feeding, the ABX + PEDV group received PEDV infection after 7 d of antibiotic feeding and further PBS feeding.

### **q-** PCR experiment

According to the manufacturer’s instructions, total RNA was extracted, and 1 mg of RNA was reversed into cDNA by reverse transcriptase ( Promega), which reverse transcribed the Moloney mouse leukemia virus. In the real-time quantitative PCR system of Applied Biosystems 7500, quantitative PCR was performed using SYBR Green mixture (TakBR Green). In the real-time quantitative PCR system of Applied Biosystems 7500, quantitative PCR was performed using SYBR Green mixture (Takara Bio). The average mRNA fold changes were calculated by 2^-ΔΔCT^, and all primer sequences are shown in Supplementary Data 1.

### Cell experiments

We stained the total LPL extracted from the lamina propria with antibodies Live/Dead, CD45, Lin1 (CD3ε, CD21, γδ T, CD11c, CD172a), and Lin2 (CD4, CD8, CD163, CD11b) and then selected L/D-CD+Lin1-Lin2-cells by flow sorting or magnetic bead sorting (STEMCELL EasySep™ PE Positive Selection Kit). CD163, CD11b) and then selected L/D-CD45+Lin1-Lin2-cells by flow sorting or magnetic bead sorting (STEMCELL EasySep™ PE Positive Selection Kit II # 17684). After obtaining, they were cultured in vitro and then flow assayed. In order to maintain the cellular activity of ILCs, for all immunological cell cultures longer than 24h, we used OP9 cells co-cultured with immune cells to enhance the survival of ILC3 cells.

### Organoid experiment

1. Piglets were euthanized by removing 8 cm of jejunum, removing the mesentery, dissecting the intestinal segments longitudinally, washing them with cold PBS until they were cleaned, cutting the segments into 0.5 cm pieces, washing them with cold PBS by blowing gently, and adding 15 mL PBS(2 mM EDTA). Let it stand at room temperature for 40 min.
2. Discard the supernatant, add 10 mL of DPBS, and blow 2 times. The supernatant was collected in a 50 mL centrifuge tube through a 70 μm cell sieve, labeled #1, and repeated 4 times. The filtrate of number 3 and 4 was centrifuged at 300 × g for 5 min, and the supernatant was discarded. The supernatant was resuspended with 1 mL of DME/F12(1% penicillin and streptomycin), transferred to a 1.5 mL centrifuge tube, centrifuged at 200 × g for 3 min, and the supernatant was discarded.
3. 250 μL of complete medium and 250 μL of Matrigel (operated on ice) were added to the precipitate and blown to mix. Pipette 50 μL in the center of a 24-well plate and place in the incubator for 30 min. Then, 500 μL medium (STEMCELL #6000) was added to each well and 500 μL PBS to the remaining wells.
4. When the organoids start to germinate, they should be passaged. First, discard the old medium and add 2 mL DME / F12, blow it up and down, and recycle it into the centrifuge tube. After centrifugation, discard the supernatant and reintroduce it into the complete medium and Matrigel.

### Isolation and Characterization of EVs

EVs were isolated from *LGG* using ultracentrifugation. The size distribution and morphology of EVs were analyzed by nanoparticle tracking analysis (NTA) and transmission electron microscopy (TEM), respectively.

### Statistics

Statistical analyses were performed using the statistical computer package, GraphPad Prism version 6, (GraphPad Software Inc., San Diego, CA). Results are expressed as means ± SEM. Statistical comparisons were made using two-way analysis of variance with Tukey’s post hoc test or Statistical comparisons were made using two-way analysis of variance with Tukey’s post hoc test or Student t test, where appropriate. Differences were considered to be significant at P < 0.05. Significance is noted as *P < 0.05, **P < 0.01, ***P < 0.001 and ****P < 0.0001among groups

### Results

#### 1. PEDV infection affects changes in the intestinal flora and ILC3 in piglets

In our previous work, we found a large number of immune cells in the piglet intestine (Wang et al., 2024), and through flow cytometric sorting and single-cell sequencing (scRNA-seq), we found that in Lin^-^ cells, there were a large number of ILCs cells (Fig. 1A), in which ILC3 expressing the signature transcription factor RORC was absolutely dominant (Fig. 1B), and also expressing, IL7R, AhR and other surface molecules, and secreted corresponding cytokines such as IL22 and IL17. The expression of these genes is important for the regulation and protection of intestinal homeostasis (Fig. 1C).

**Fig. 1.**
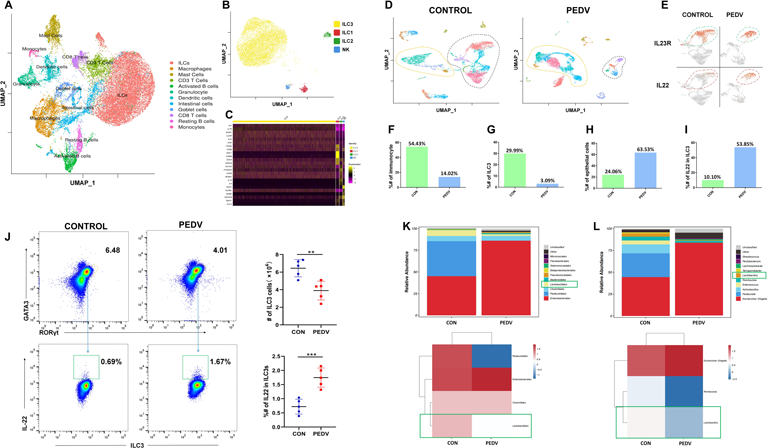
PEDV infection resulted in abnormal changes in both the intestinal flora and ILC of piglets. A: UMAP plots showing the immune landscape of the 12 subpopulations of the intestine cell(Selection by flow cytometry). B: UMAP plots showing the immune landscape of the 4 clusters of intestine cells (ILC regrouping). Cells are color-coded according to the defined subset (ILC3: yellow; ILC1: red; ILC2: green; NK: blue). C: Heatmap showing the marker genes with some highest differentially expressed in each cluster, Abscissa shows different clusters, and ordinate shows differentially expressed gene names. D: UMAP plots showing the immune landscape of intestine cells in the jejunum of piglets in both PEDV-infected and normal states. E: The amount of ILC3 significantly decreased, while the secretion of IL-22 was significantly increased. F: Variations in the quantity of immune cells. G: Variations in the quantity of ILC3. H: Variations in the quantity of epithelial cells. I: Differential changes in the secretion of IL-22 by ILC3. J: The amount of ILC3 in the intestinal lamina propria of PEDV-infected piglets significantly decreased, and the secretion of IL-22 increased. K: Changes in Lactobacillales orders in the intestinal tract of piglets. J: Changes in Lactobacillus Genus in the intestinal tract of piglets.

Through scRNA-seq analysis, we found that PEDV infection in piglets led to a significant decrease in the number of immune cells in the jejunum, including T cells and ILC3 (Fig. 1D, F, G). In contrast, non-immune cells, such as epithelial cells, exhibited abundant proliferation during the infection. (Fig. 1D, H). Despite the significant decrease in the number of ILC3 cells in the jejunum, the secretion of IL-22 was markedly increased (Fig. 1E, I). These results suggest that the increased secretion of IL-22 by ILC3 cells may play a crucial role in conferring resistance against PEDV during PEDV infection. Our flow cytometry assay results were consistent with the single-cell results, the flow gating strategy in Fig. S1A. These results also confirmed a significant decrease in the number of ILC3 cells in the intestinal lamina propria of PEDV-infected piglets, accompanied by an up-regulation of IL-22 secretion by ILC3 cells (Fig. 1J). Furthermore, our findings revealed significant alterations in the flora of the jejunum in PEDV-infected piglets. At both the order level and the genus level, there was a notable decrease in the levels of *Lactobacillales* orders and *Lactobacillus* genus (Fig. 1K, L).

To further investigate the underlying cause of the significant changes in ILC3 during PEDV infection, we aimed to discern whether these changes were primarily driven by viral infection or influenced by alterations in the bacterial flora. To explore this, we conducted an experiment involving antibiotic treatment to manipulate the structure of the flora. By observing the effects of this flora change on the body’s immunity, we aimed to gain insights into the intricate relationship between the microbial composition and the immune response.

#### 2. Alteration of intestinal flora may affect ILC3 development

In our study, we initially established a dysregulation model of piglets’ microflora. We observed significant impacts on the immune development of piglets with dysflora. Specifically, there was a noticeable reduction in the number of CD4-T cells in the jejunum lamina propria (Fig. S2A). Additionally, the development of ILC3 was even more severely affected, with a significant decrease in cell numbers and an increase in IL-22 secretion (Fig. 2A). Meanwhile, antibiotic treatment resulted in the decrease of flora richness and *lactobacillus* number in the jejunum of piglets (Fig. 2B, C). Linear discriminant analysis Effect Size (LEfSe) analysis identified significantly different taxa between groups, which were significantly enriched from the point of view of order, genus, and species, respectively, belonging to *Lactobacillales*, *Lactobacillaceae*, and *Lactobacillus* (Fig. 2D, E, F, G).

**Fig. 2.**
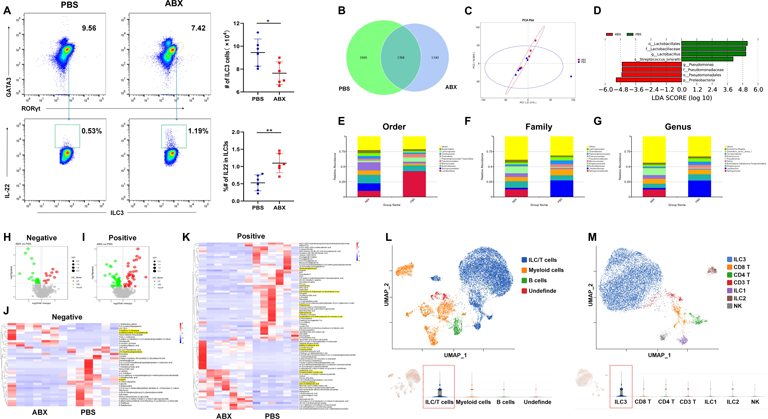
The imbalance of intestinal flora may affect the development of ILC3. A: The number of ILC3 in the intestinal lamina propria of piglets with intestinal flora disturbance decreased significantly, and the secretion of IL-22 increased. B: The Venn diagram showed the difference of intestinal flora species between the PBS and ABX groups (α Diversity). C: PCA analysis between the PBS and ABX groups (β Diversity). D: LDA Effect Size (LEfSe) analysis between the PBS and ABX groups (β Diversity). E: Changes in Lactobacillales orders in the intestinal tract of piglets. F: Changes in Lactobacillaceae family in the intestinal tract of piglets. G: Changes in Lactobacillus genus in the intestinal tract of piglets. H: Changes in differential metabolites in the negative ion mode. I: Changes in differential metabolites in the positive ion mode. J: Different substances were observed between the groups in negative ion mode, with a particular focus on substances that are associated with AHR. K: Different substances were observed between the groups in negative ion mode, with a particular focus on substances that are associated with AHR. L: UMAP and violin plots showing the AHR was prominently expressed in immune cells within pig intestines. M: UMAP and violin plots showing the AHR was prominently expressed in ILC3 within pig intestines.

The above results suggest that the number of immune cells, especially ILC3, was significantly affected under both PEDV infection and antibiotic treatment conditions and that this change persisted when only the flora was changed. The flora may play a very important role in the interactions with ILC3, and if there is a regulatory effect of the flora on ILC3, by what pathway does the flora regulate ILC3? Up-regulation of indole analogs, including bisindolylmaleimide I and indoxyl sulfate, has been reported in PEDV infection (Li et al., 2022). Non-targeted metabolomics was further analyzed for the metabolites in jejunum samples. As a result, we found significant changes in some AhR-related regulatory substances in both positive and negative ion modes (Fig. 2H, I, J, K), especially many indoles that have been repeatedly shown to have a significant activating effect on the AhR. We further performed enrichment analyses and found a significant enrichment in important immune-regulatory pathways such as tryptophan. To investigate whether differential microbial communities produce differential metabolites, we conducted a combined analysis of 16S and metabolomics. The results revealed a significant decrease in the abundance of regulatory substances secreted by probiotic bacteria, leading to a reduction in their overall levels. However, unfortunately, we did not directly associate the metabolites with the production by *Lactobacillus* (Figure S2B, C).

So far, the expression of the AhR receptor has been studied in immune cells in humans and mice, but its expression has not been reported in pigs. Furthermore, the expression of AhR in immune cells in the intestine is unknown. We found that immune cells in the porcine intestine highly expressed AhR receptors by single-cell analysis (Fig. 2L), and further analysis revealed that ILC3 in the intestine expressed high levels of AhR (Fig. 2M). In human intestinal ILC, both ILC3 and ILC1 highly expressed AhR receptors (Fig. S2D), whereas only ILC2 expressed AhR in the murine upper intestine (Fig. S2E). These results suggest that porcine and human intestinal ILC3 exhibit high levels of AhR activation, indicating that ILC3 in the intestine may respond significantly to stimulation by AhR ligand-related substances.

#### 3. Feeding *LGG* can enhance the piglets’ immunity against PEDV infection

To explore whether *Lactobacillus* can resist PEDV infection in the gut, we selected the standard *Lactobacillus* strain *LGG* to supplement piglets with an intestinal flora imbalance and then carried out a PEDV challenge experiment. Our objective was to determine whether feeding *LGG* could enhance intestinal immunity, specifically by activating ILC3 cells, thereby promoting intestinal homeostasis and conferring resistance against PEDV infection.

It was found that *LGG* supplementation significantly reduced the pathological changes of PEDV infection in piglets, and the intestinal tract was healthy in the CON group. The villi of the jejunum of piglets in the ABX+*LGG*+PEDV group were shortened and atrophied, and the epithelial cells were mildly diseased. The villi of the jejunum in piglets from the ABX+PEDV group exhibited shortened, fragmented, and broken structures. Additionally, the intestinal epithelial cells underwent typical histopathological changes such as vacuolization, fusion lesions, necrosis, and detachment following PEDV infection (Fig. 3A). The ratio of villus height to crypt depth (VH/CD) of the jejunum villi of piglets in each infection group was found to be significantly lower in the ABX+PEDV group than in the ABX+*LGG*+PEDV group (Fig. 3B). The diarrhea scoring results proved that feeding *LGG* alleviated diarrhea in piglets (Fig. 3C). Next, we found significant differences in viral loads in the duodenum, jejunum and ileum between the ABX+*LGG*+PEDV group and the ABX+PEDV group, especially in the jejunum and ileum, where the viral copy number was significantly decreased after feeding *LGG* (Fig. 3E).

**Fig. 3.**
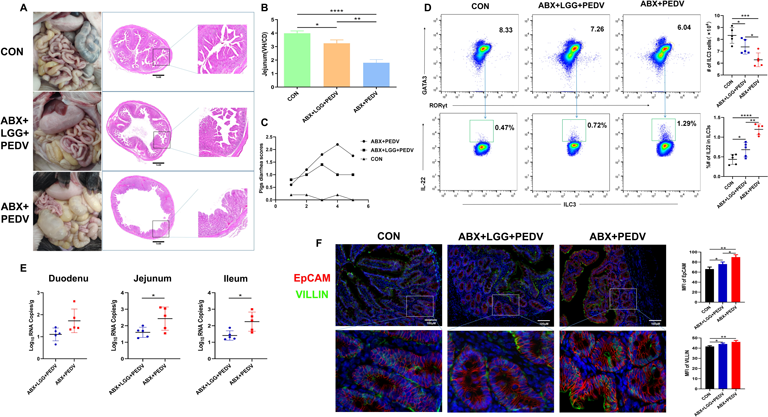
LGG can help piglets resist PEDV infection. A: Feeding LGG can significantly alleviate the pathological changes of jejunum in piglets. B: The ratio of villus height to crypt depth (VH/CD) of intestinal villi of piglets. C: Piglet diarrhea score. 0= normal stool, 1= soft but formed stool, 2= semi-liquid stool, 3= watery diarrhea, with a score of 2 or more considered diarrhea. D: Flow cytometry showed the change of ILC3 quantity and IL-22 secretion in the intestinal tract of piglets. E. The q-PCR results showing viral load in the intestines of piglets. F: Immunofluorescence assay showed the expression of EpCAM and VILLIN in the gut, Analysis by image J software.

Flow results also demonstrated a significant increase in the amount of ILC3 in the intestines of the ABX+*LGG*+PEDV group, which was close to that of the CON group and significantly higher than that of the ABX+PEDV group. The secretion of IL-22 was also significantly lower than that of the ABX+PEDV group, suggesting that the inflammatory state of the jejunum intestinal tract of the piglets was ameliorated in response to the infection (Fig. 3D). A dramatic improvement also occurred in the abnormal activation state of the intestinal tract through stimulation by feeding *LGG* to promote proliferation of epithelial cells and the production of IL-22 after viral infection, including the expression of proteins such as EpCAM and VILLIN in the intestinal tract (Fig. 3F). There was also a significant reduction in the cell proliferation-related proteins LGR5 and Ki-67 (Fig. S3A).

Our results at the mRNA level were consistent with the results at the protein level, revealing that inflammation-related genes such as IL-22 and IL-1β, the epithelial gene EpCAM, the villi gene VILLIN, the proliferation and differentiation-related gene MKI-67, the ISC labeling genes Lgr5, Ascl2, Olfm4, and the Paneth cell activation genes Lyz1 and REG3b were all significantly reduced (Figure S3B). Feeding *LGG* also significantly increased the abundance of intestinal flora in piglets, particularly the number of the *Lactobacillus* genus, and elevated the content of other probiotics, such as *Streptococcus spp*, suggesting that feeding *LGG* can promote the recovery of the intestinal flora (Fig. S3C).

#### 4. *In vitro* cellular experiments demonstrated that *LGG* metabolites promote ILC3 cell activation to resist PEDV

We first successfully constructed an in vitro culture model of ILC3 (Fig. S4A), performed a flow cytometry assay under different stimulant-inducing conditions, and developed a gating strategy (Fig. S4B). We found that both *LGG*-secreted extracellular vesicles (EVs) and Indole-3-carbinol (I3C) could significantly promote ILC3 proliferation and secretion of IL-22 through activation of the AhR. The effect disappeared when the AhR antagonist CH-223191 (CH) was added (Fig. 4A).

**Fig. 4.**
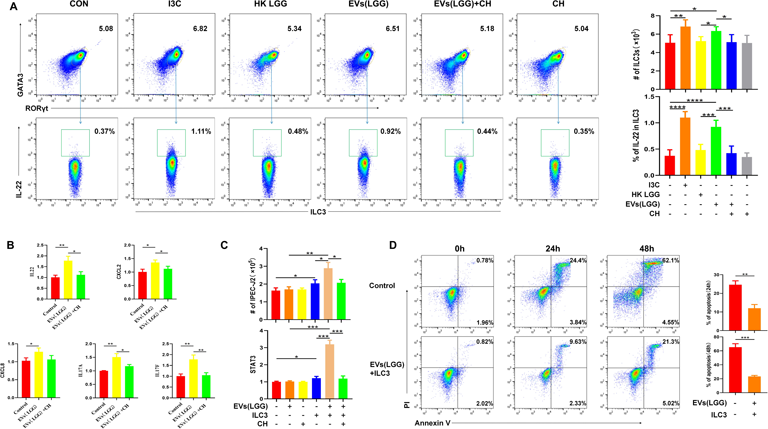
Experimental results of LGG metabolites promoting ILC3 cell activation. A: Flow cytometry showed that I3C and EVs can promote the amount of ILC3 and the secretion of IL-22 under co-culture conditions. B: Transcriptional levels of ILC3-related cytokines stimulated by LGG metabolites were measured by q-PCR. C: Statistics of the number of IPEC-J2 cells; The transcription level of STAT3 gene in IPEC-J2 cell species was detected by q-PCR. D: Flow cytometry calculates the percentage of cells that die. The Annexin V-/PI+ cells in the upper left quadrant may represent cell fragments lacking cell membranes or dead cells resulting from other causes. The Annexin V-/PI-cells in the lower left quadrant are considered normal and alive. The Annexin V+/PI+ cells in the upper right quadrant are indicative of late apoptotic cells. Lastly, the Annexin V+/PI-cells in the lower right quadrant also correspond to early apoptotic cells.

Previous correlational studies have indicated the possibility of ILC transformation within the organism, particularly when induced by specific conditions (Bal et al., 2020). To investigate whether the change in the number of ILC3 is due to the transformation of other immune cells, we conducted magnetic bead sorting on ILCs cells and subsequently examined them using SFSE staining(Fig. S4C). We discovered that ILC3 is the primary cell population within ILCs where proliferation occurs. The proliferation of ILCs cells dominated by ILC3 was significantly stronger than that of other non-ILC cells (Fig. S4D). The expression of ILC3-related cytokines, including IL-22, IL-17A, IL-17B, CXCL2, and CXCL8, were all found to be significantly up-regulated (Fig. 4B).

Our study investigated whether the activation of ILC3 and secretion of IL-22 promoted by EVs are effective in enhancing the resistance of porcine epithelial cells to PEDV infection. We established a co-culture model of ILC3 and IPEC-J2 cells (Fig. S4E) and observed that IL-22 secreted by EVs-promoted ILC3 had a positive impact on IPEC-J2 cell proliferation and the activation of STAT3 (Fig. 4C). Furthermore, when EVs and ILC3 were added to co-cultured IPEC-J2 cells after PEDV infection, it significantly influenced the outcome of PEDV infection by preventing apoptosis of IPEC-J2 cells (Fig. 4D), the gating strategy in Fig. S4F.

These results suggest that EVs derived from *LGG* can stimulate the proliferation and secretion of IL-22 by ILC3 through the AhR receptor on ILC3. This process promotes the proliferation and activation of epithelial cells and enhances the expression of the STAT3 gene. Therefore, it alleviates IPEC-J2 apoptosis caused by PEDV infection.

#### 5. Porcine intestinal organoid proving experiments

We successfully constructed an in vitro model of organoids from the jejunum of piglets. The size of isolated intestinal crypts was about 15 μm, and they were cultured with matrigel. The first generation of organoids grew slowly and developed into mature bodies with a diameter of about 200 μm by the eighth day and were continued to be cultured. Then they died and fragmented, and displayed a large number of buds in 5-8 d and could be propagated through passaging culture(Fig. 5A). Subsequently, the cultured organoids grew rapidly after passaging, reaching a size of 100 μm in three d and matured into bodies with a diameter of 200 μm within 5 d (Figure S5B).

**Fig. 5.**
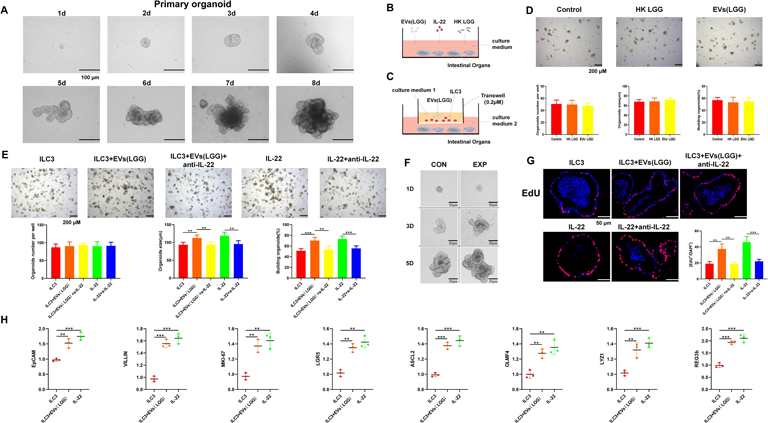
Piglet intestinal organoids experiment. A: Crypts isolated from piglet intestines were cultured in vitro, and the images depict the changes in organoid size over an eight-day period. B: HK LGG and EVs were added to the intestinal organoid culture hole. C: The co-culture system consisted of intestinal organoids, along with associated stimulatory molecules and ILC3 cells. D: Effects of HK LGG and EVs on the growth of intestinal organoids. Including size of organoids treated with/without HK LGG and EVs, total organoids number and budding organoids percentage of total organoids per well (day 3) (n=50). E. The effects of adding ILC3, ILC3+EVs, ILC3+EVs(LGG)+anti-IL-22, IL-22, IL-22+anti-IL-22 in organoid medium on organoid growth were statistically analyzed (n=50). F: Organoid growth of the experimental group and the control group began to show significant differences on day 2. G: Organoids were stained with EdU (red). Nuclei are stained blue. EdU-positive cells were found in the transit-amplifying region with obvious differences in the percentages among samples (n=10). H: Transcriptional changes of EpCAM, VILLIN, MKI-67, Lgr5, Ascl2, Olfm4, Lyz1 and REG3b genes were detected by q-PCR (n=3).

Previous experiments have shown that EVs can stimulate the secretion of IL-22 from ILC3. Therefore, it is worth investigating if IL-22, secreted by ILC3, can promote the growth of organoids. In our study, we initially supplemented the intestinal organoid medium with HK *LGG* and EVs derived from *LGG* (Fig. 5B). We then monitored the number of organoids, organoid size, and germination rate throughout the organoid growth process. Interestingly, we observed that the addition of HK *LGG* and EVs alone did not have any noticeable effect on the development of organoids (Fig. 5D).

In the next phase of our study, we successfully established a co-culture system involving intestinal organoids, stimulatory molecules, and ILC3 cells (Fig. 5C). Initially, we attempted to culture ILC3 cells by adding them directly to the matrigel. However, this method proved unsuitable for the survival of ILC3 cells and resulted in severe cell death and fragmentation within 24 h. To overcome this issue, we performed co-cultures using a transwell system with a pore size of 0.4 μm. We renewed the upper layer of ILC3 cells every 24 h. Notably, we observed that the growth of the organoids was significantly enhanced in the co-culture system when EVs+ILC3 and IL-22 were added. Within the first two d, the organoids experienced rapid growth, reaching a size of 70 μm. Subsequently, they started to exhibit significant budding and divided into multiple crypts on the third day. The organoids continued to grow rapidly, whereas the promotional effect of EVs and IL-22 on organoid growth disappeared upon the addition of an antibody specific to IL-22. Importantly, the number of organoids was not affected (Fig. 5E). Comparing the experimental group (EVs+ILC3, IL-22) with the control group (including ILC3, ILC3+EVs(*LGG*)+anti-IL-22, IL-22+anti-IL-22), we found that the experimental group significantly promoted the growth of the organoids. Notably, a significant difference was observed on the third day during germination, and by the sixth day, the size of the organoids in the experimental group showed significant differences compared to the control (Fig. 5F).

Next, immunofluorescence results revealed significant expression of proliferation-related proteins in organoids after addition of co-cultures of EVs+ILC3 and IL-22, including increased expression of 5-Ethynyl-2′-deoxyuridine (EdU) (Fig. 5G), the ISCs activation-related protein LGR5, and the cell proliferation-related protein Ki67; a significant increase in expression of the epithelial Significantly increased expression of the epithelial protein EpCAM and the chorionic protein VILLIN was also found (Fig. S5C). Q-PCR results also identified the epithelial gene EpCAM, the chorionic gene VILLIN, the proliferation and differentiation-associated gene MKI-67, and the ISCs marker genes Lgr5, Ascl2, and Olfm4 in the class of organoids following the incorporation of co-cultures of EVs+ILC3 and IL-22; and Paneth cell activation genes Lyz1 and REG3b were significantly increased (Fig. 5H).

These results demonstrated that the EVs of LGG could significantly promote the growth and development of organoids and activate ISCs, Paneth cells, epithelial cells, and other cells in organoids.

#### 6. The metabolites of *LGG* are rich in AhR ligands, such as indole compounds

We further examined the *LGG*-produced EVs and found the diameter of the EVs to be around 70 nm by nanoflow cytometry (Fig. 6A). The signature horseshoe-shaped structure of the EVs, which contained a large amount of *LGG* metabolites, was visualized under an electron microscope (Fig. 6B).

**Fig. 6.**
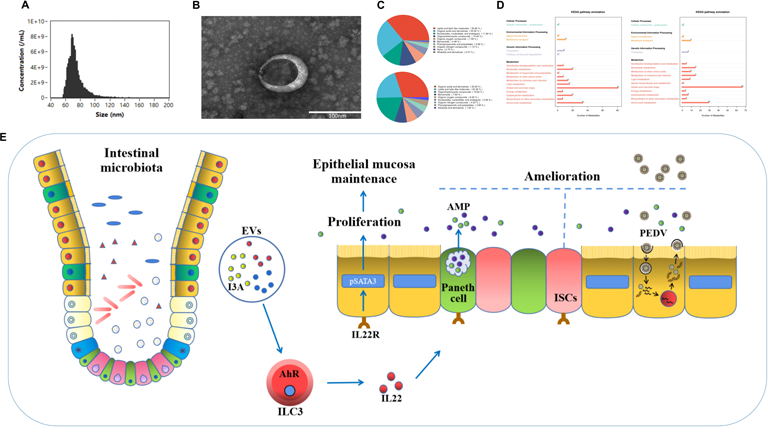
The detection of EVs (LGG) and the pathway of maintaining intestinal immune homeostasis. A: Results of nanoflow cytometry detection of EVs (LGG). B: Observation of EVs (LGG) under electron microscope. C: Detection of EVs (LGG), classification of LGG metabolites in positive and negative ion modes. D: KEGG enrichment analysis was conducted for LGG metabolites. E: The pathway of EVs (LGG) to maintain intestinal immune homeostasis in piglets.

We enriched and collected the extracellular vesicles (EVs) through ultrafast centrifugation. Subsequently, we conducted off-target metabolism detection. In total, 414 substances were identified in the positive mode (POS), while 326 substances were detected in the negative mode (NEG). We observed a significant presence of lipids and lipid-like molecules, as well as organic acids and derivatives, in both the POS and NEG modes (Fig. 6C). We found a wide range of substances that can interact with AhR, including indole compounds such as L-5-Hydroxytryptophan and Indole-3-acrylic acid, as well as several ketone compounds, and some ketones. Research has indicated that, apart from indoles, ketones can also interact with AhR (Amakura et al., 2008).

We conducted KEGG enrichment analysis on the total metabolites and found significant enrichment in the Metabolism pathway. Additionally, enrichment was observed in pathways such as Cellular Processes, Environmental Information Processing, and Genetic Information Processing (Fig. 6D).

Our study provides insights into the interaction between ILC3s in the porcine small intestine and *Lactobacillus*, aiming to understand better how they interact. We have demonstrated that oral *LGG* is a potential approach for preventing PEDV infection in pigs. It can alleviate intestinal inflammation, intestinal damage, and clinical diarrhea symptoms caused by PEDV infection. We found that EVs (*LGG*) can activate ILC3s and promote IL-22 secretion, thereby influencing intestinal stem cell regeneration and epithelial protection. Ultimately, this helps to purify the gut environment and resist PEDV infection. In summary, our research demonstrates that oral *LGG* promises to be a potential method for preventing PEDV infection in pigs. Furthermore, it opens up new avenues for preventing PEDV infection in piglets.

## Discussion

Recently, we discovered the presence of ILCs in the jejunum of piglets. These cells are predominantly ILC3 and play a crucial role in maintaining intestinal immune homeostasis. In our study, we further elucidated the interaction between ILC3 and the gut microbiota, particularly *Lactobacillus*, in the jejunum of piglets to maintain intestinal homeostasis. Based on our findings and existing literature, we constructed a pathway diagram illustrating how EVs produced by *LGG* facilitate the secretion of IL-22 by ILC3 cells. IL-22, in turn, acts on downstream targets such as epithelial cells, Paneth cells, and ISCs to regulate intestinal immune function, thereby conferring protection against PEDV infection (Figure 6E).

The gut microbiota plays a crucial role in the development and maintenance of the host immune system, and its complexity is undeniable. It stabilizes mucosal function by maintaining the integrity of the gut barrier and balances the inflammatory response with immune tolerance through the induction of T lymphocyte differentiation. Additionally, the gut microbiota promotes the development of early B lymphocytes in the lamina propria of the mouse small intestine (Zegarra-Ruiz et al., 2021; Rooks and Garrett, 2016). At the same time, the immune system also regulates the microbial community in a certain way to maintain relative balance. However, changes in this balance are often associated with the occurrence or progression of diseases. Probiotics have a positive effect on the gut microbiota, especially in the innate and adaptive immune systems. Preclinical studies and clinical practice have shown that the use of probiotics can limit the overgrowth of pathogenic bacteria and control the host’s pathological processes (Pitocco et al., 2020). Furthermore, the interaction between bacteria and host cells is a complex process that has yet to be fully understood and deserves further exploration. Our research revealed significant changes in the gut microbiota during PEDV infection, including a decrease in the abundance of beneficial bacteria such as *Lactobacillus*. However, oral administration of *LGG* can greatly alleviate dysbiosis and increase the diversity of the gut microbiota.

AhR is a widely expressed transcription factor in immune system cells, particularly playing a crucial role in intestinal immune cells. Studies have shown that AhR activation specifically alters innate and adaptive immune responses and participates in the regulation of cell differentiation and inflammation-related gene expression. Furthermore, inducible AhR activation may serve as a mechanism by which the gut microbiota promotes mucosal homeostasis. In comparison to its role in intestinal immune cells, the function of AhR in intestinal epithelial cells (IECs) has not been extensively studied. Previous research has found a close association between AhR in IECs and the maintenance of epithelial barrier function (Qian et al., 2022). One of the endogenous ligands for AhR is a metabolite of tryptophan. The gut microbiota can convert tryptophan into various molecules, including ligands for AhR, such as indole and its derivatives (e.g., IAld, IAA, IPA, IAAld, and indole-3-acrylic acid). The AhR signaling pathway is an essential component for maintaining barrier immune responses, promoting epithelial cell recovery, preserving barrier integrity, and regulating certain immune cell functions to maintain gut homeostasis. The gut microbiota and tryptophan and its metabolites interact closely in various aspects, including gut barrier function, gut immunity and endocrine activity, and intestinal motility (Su et al., 2022). Current research has described a few commensal bacteria, such as *Lactobacillus*, capable of producing AhR ligands (Agus et al., 2018). However, the significance of these complex phenomena in the intestinal system still requires further investigation. One study discovered that adaptive lactobacilli can expand and produce an AhR ligand, I3A, which contributes to the transcription of IL-22 dependent on AhR. Therefore, the microbiota-AhR axis may serve as an important strategy for modulating host mucosal immune responses in symbiotic evolution (Zelante et al., 2013).

Additionally, lactobacilli can regulate immune cells through AhR by producing substances like tryptophan and expressing tryptophanase, thus promoting the production of indole-3-propionic acid associated with human health (Liu et al., 2023). Our research findings demonstrate that in a healthy state of piglet gut microbiota, there is a higher proportion of *Lactobacillus*. Additionally, we observed a significant presence of substances in the gut that interact with AhR. Moreover, AhR exhibits notable upregulation of porcine immune cells, particularly ILC3, indicating a potentially tighter regulatory relationship between ILC3 and the gut microbiota via AhR in the pig gut. In states of PEDV infection or dysbiosis, the levels of substances interacting with AhR significantly increase in the gut. Activation of ILC3 and secretion of IL-22 are subsequently promoted through AhR, thereby exacerbating inflammation and accelerating stimulation and activation of downstream receptor cells by IL-22. However, oral administration of *LGG* to piglets leads to significant restoration of gut microbiota balance and the regulatory relationship between metabolites and AhR.

IL-22 is a member of the IL-10 family that can be produced by ILCs and CD4-T cells. The IL-22-IL-22R signaling axis plays a crucial role in integrating immune responses with mucosal surface barrier function (Makowski et al., 2020; Dudakov et al., 2015). Increasing evidence suggests that IL-22 plays an important role in inflammatory bowel disease (IBD) (Mizoguchi 2012). IL-22 produced by ILC3 is essential for maintaining intestinal homeostasis and provides early protection to epithelial barrier function during inflammation and injury (Zenewicz et al., 2008). The interaction between the microbiota and IL-22 is central to regulating the barrier sites in intestinal homeostasis. On the one hand, IL-22 promotes intestinal barrier function by inducing antimicrobial peptides, mucins, and other beneficial factors from epithelial cells, thereby modulating the composition of the gut microbiota. On the other hand, the gut microbiota also regulates the production of intestinal IL-22, although the underlying mechanisms are not fully understood. Research has reported that many functions of the gut microbiota in regulating health and disease are mediated through their metabolites (Lynch and Pedersen 2016, Rooks and Garrett 2016). In our in vitro experiments using IPEC-J2 cells and piglet intestinal organoids, we found that EVs produced by *LGG* can promote the activation of ILC3 cells and the production of IL-22. The generated IL-22 can act on IPEC-J2 cells, promoting their proliferation, activating the STAT3 signaling pathway, and conferring resistance against PEDV infection. Moreover, the produced IL-22 can also act on Paneth cells, ISCs, and epithelial cells within the organoids to promote their growth and development.

Investigators have found that IFN-γ and IL-22 act synergistically to induce interferon-stimulated genes and control rotavirus infection. Therefore, this pathway may not be specific to PEDV infection and could potentially play a role in other diseases as well (Hernández et al., 2015). Although we observed significant changes in the secretion of ILC3 and IL-22 in the jejunum of piglets following exposure to *Lactobacillus*, it is still unknown whether other bacteria can significantly stimulate ILC3 to secrete IL-22. Could it be possible that the balance between lactobacilli and ILC3 is disrupted, leading to dysregulation of the existing regulatory relationship and a significant increase in the number of harmful bacteria, thereby further stimulating ILC3 activation? These questions remain unanswered, and we hope to address them in future research.

Currently, there is increasing interest in studying the interactions between microorganisms and the immune system. On the one hand, the immune system can regulate and shape the microbial community, and on the other hand, the established microbial community can promote the development of the host’s immune system and provide signals for subsequent immune responses. However, our understanding of the interactions between microorganisms and the immune system is still limited, and unraveling these complexities requires interdisciplinary collaboration and innovation. Numerous intrinsic factors influence the balance within the animal body. Although our research focuses on the interaction between intestinal lactobacilli and ILC3 through the AhR, it is important to note that there are countless regulatory pathways within the animal body. Moreover, research on pigs is relatively scarce, and much of our knowledge comes from studies conducted on humans and mice. Therefore, we cannot fully comprehend the complete role of ILC cells in the intestinal immunity of pigs. Furthermore, considering that ILC3 is an emerging cell type in pigs, our work is just the beginning. In the future, we will explore the changes in this cell population during intestinal immunity and disease states, laying the foundation for studying intestinal immunity in pigs.

## Acknowledgments

We thank Dr. Guangliang Liu Provided to us with the PEDV strain. We thank Dr. Xuechao Gao from Geneu Biotech Co., Ltd., who helped us obtain monoclonal antibodies against porcine IL-22. We thank Novogene Co., for providing technical services such as detecting and analyzing of scRNA-seq raw data, 16S rDNA sequencing and Metabolome sequencing.

## Data availability statement

The raw data for this article were deposited in the National Center for Biotechnology Information (NCBI) Sequence Read Archive (SRA) database, Gene Expression Omnibus (GEO) database and EBI database. The source of the jejunal single cell data after PEDV infection in piglets is GSE175411. Single cell data of piglet jejunum ILCs in PRJNA907920. We comparisons with Human intestinal ILCs (GSA-Human: HRA000919) and mouse intestinal ILCs (GSE166266). 16S rRNA amplicon sequencing data in PRJNA526581, PRJNA1048561 and PRJNA1048648.

## Author contributions

Cell isolation, J.H.W., Y.B.Z., and T.C.; data analysis, J.H.W., H.Y.B., M.G., Y.S., and Y.Y.L.; manuscript preparation and writing, J.H.W., Y.Y.L., M.Y.C., and H.Y.B.; Information collection, Y.B.Z., T.C., and H.Y.B.; supervision and project administration, C.F.W., Y.Z., and X.C. Preparation of experimental reagent materials, C.W.S., J.Y.G., J.Z.W., N.W., W.T.Y., Y.L.J., H.B.H., D.Z., J.T.H., G.L.Y. All authors contributed to the article and approved the submitted version.

## Ethics declarations

### Consent to publish

All authors have approved the content of this manuscript and provided consent for publication.

### Conflict of Interest

The authors declare that the research was conducted without any commercial or financial relationships that could be construed as a potential conflict of interest.

### Ethics

The animal management procedures and all laboratory procedures abided by the regulations of the Animal Care and Ethics Committees of Jilin Agriculture University, China.

### Funding

This work was supported by the National Natural Science Foundation of China ( 32273043, 32202890, U21A20261), the Science and Technology Development Program of Changchun City (21ZY42), the Science and Technology Development Program of Jilin Province(20200402041NC), and China Agriculture Research System of MOF and MARA (CARS-35).

This paper does not report the original code. Any additional information required to reanalyze the data reported in this paper is available from the lead contact upon request.

---

**Fig. S1.**
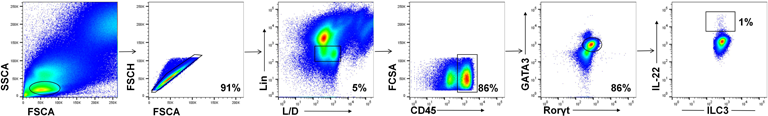
A: The 903 gating of ILC subsets.

**Fig. S2.**
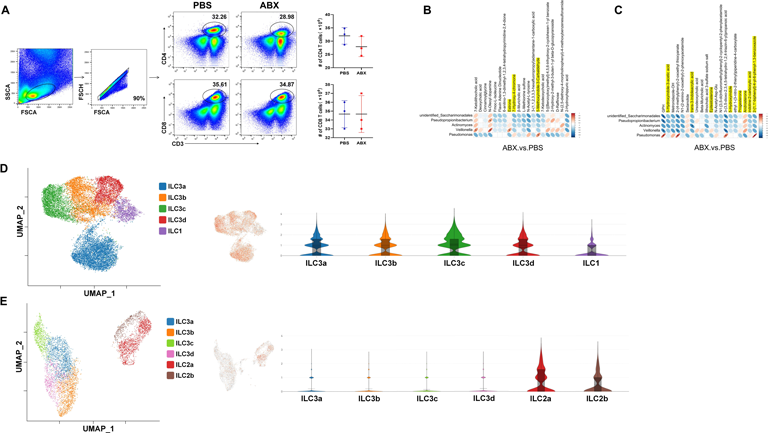
A: The statistical analysis of flow cytometry results revealed that antibiotic treatment had an impact on CD4-T cell in LPL, although the difference was not significant. However, it did not have any effect on CD8-T cell. B/C: We showed the results of the correlation analysis between differential metabolites and differential microbial communities, including the positive and negative ion modes. D: UMAP and violin plots showing the AHR was prominently expressed in ILC3 and ILC1 within human intestines. E: UMAP and violin plots showing the AHR was prominently expressed in ILC2 within mice intestines.

**Fig. S3.**
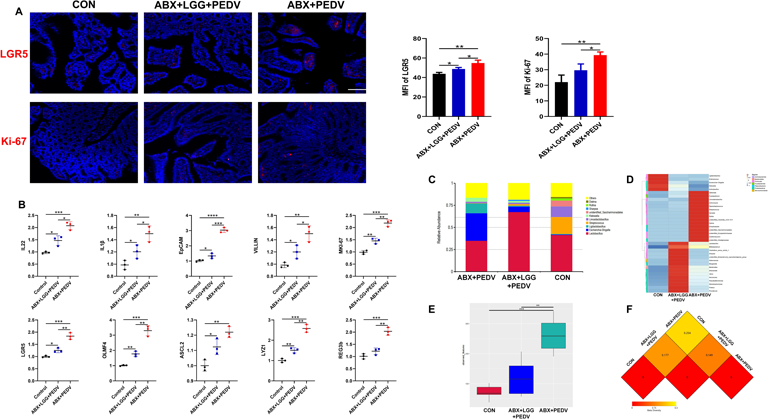
A: Immunofluorescence assay showed the expression of LGR5 and Ki-67 in the gut, Analysis by image J software. B: q-PCR experiment counted the secretion of related cytokines in the intestines of piglets in each group. C: Bar chart analysis of the genus-level microbiota in the jejunum of piglets from different groups. D: Heatmap depicting the differences in genus-level microbiota composition in the jejunum of piglets from different groups. E: Alpha diversity refers to the analysis of differences between groups, and box plots can visually represent the median, dispersion, maximum value, minimum value, and outliers of species diversity within each group. F: In the study of beta diversity, four metrics, namely Weighted unifrac distance, Unweighted unifrac distance, Jaccard distance, and Bray-Curtis distance, are used to sure the dissimilarity coefficient between two samples. A smaller 924 value indicates that the two samples have less differences in terms of species diversity.

**Fig. S4.**
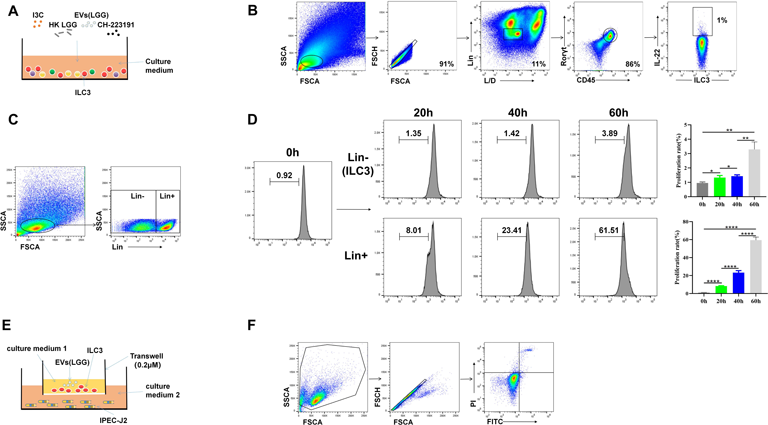
A: Figure depicting an in vitro co-culture model of ILC3. B: Schematic representation of the flow cytometry gating strategy. C: Illustration of magnetic bead sorting demonstrated by flow cytometry results. D: Proliferation assay of ILC cells, quantitatively analyzed using flow cytometry. E: Schematic diagram of an in vitro co-culture model of ILC3 with IPEC-J2. F: Flow cytometry gating strategy.

**Fig. S5.**
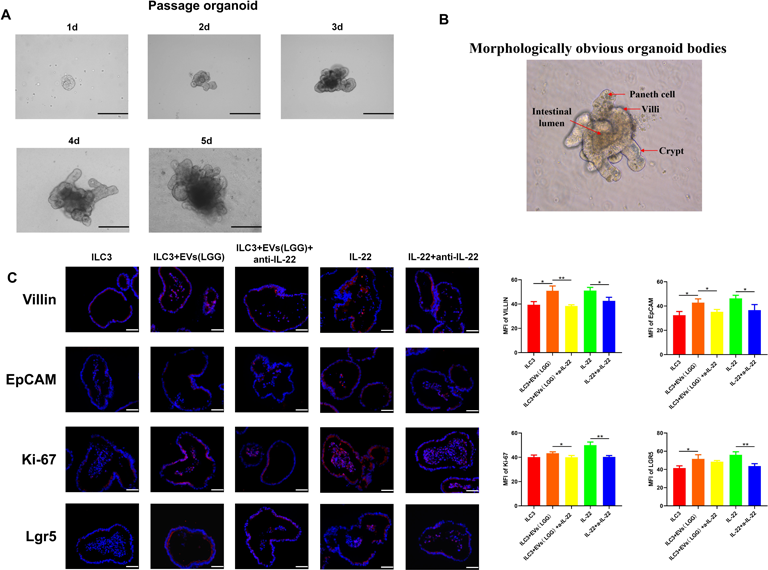
A: Growth dynamics of porcine intestinal organoids during 5 days of serial passage culture. B: Schematic representation of the composition of porcine intestinal organoids. C: Immunofluorescence experiment demonstrating the expression of proteins such as Villin, EpCAM, Ki-67, Lgr5, among others, between different groups (n=10).

## Notes

### Competing Interest Statement

The authors have declared no competing interest.

## References

Agus A, Planchais J, Sokol H (2018). Gut Microbiota Regulation of Tryptophan Metabolism in Health and Disease. Cell host & microbe 23: 716–724.

Amakura Y, Tsutsumi T, Sasaki K, Nakamura M, Yoshida T, Maitani T (2008). Influence of food polyphenols on aryl hydrocarbon receptor-signaling pathway estimated by in vitro bioassay. Phytochemistry 69: 3117–3130.

Aparicio-Domingo P, Romera-Hernandez M, Karrich J, Cornelissen F, Papazian N, Lindenbergh-Kortleve D et al (2015). Type 3 innate lymphoid cells maintain intestinal epithelial stem cells after tissue damage. The Journal of experimental medicine 212: 1783–1791.

Backert I, Koralov S, Wirtz S, Kitowski V, Billmeier U, Martini E et al (2014). STAT3 activation in Th17 and Th22 cells controls IL-22-mediated epithelial host defense during infectious colitis. Journal of immunology (Baltimore, Md: 1950) 193: 3779-3791.

Bal S, Golebski K, Spits H (2020). Plasticity of innate lymphoid cell subsets. Nature reviews Immunology 20: 552–565.

Chen J, Cui Y, Wang Z, Liu G (2020). Identification and characterization of PEDV infection in rat crypt epithelial cells. Veterinary microbiology 249: 108848.

Dudakov J, Hanash A, van den Brink M (2015). Interleukin-22: immunobiology and pathology. Annual review of immunology 33: 747–785.

Gury-BenAri M, Thaiss C, Serafini N, Winter D, Giladi A, Lara-Astiaso D et al (2016). The Spectrum and Regulatory Landscape of Intestinal Innate Lymphoid Cells Are Shaped by the Microbiome. Cell 166: 1231–1246.e1213.

Hanash A, Dudakov J, Hua G, O’Connor M, Young L, Singer N et al (2012). Interleukin-22 protects intestinal stem cells from immune-mediated tissue damage and regulates sensitivity to graft versus host disease. Immunity 37: 339–350.

Hernández P, Mahlakoiv T, Yang I, Schwierzeck V, Nguyen N, Guendel F et al (2015). Interferon-λ and interleukin 22 act synergistically for the induction of interferon-stimulated genes and control of rotavirus infection. Nature immunology 16: 698–707.

Jung K, Saif L (2015). Porcine epidemic diarrhea virus infection: Etiology, epidemiology, pathogenesis and immunoprophylaxis. Veterinary journal (London, England: 1997) 204: 134-143.

Kiss E, Vonarbourg C, Kopfmann S, Hobeika E, Finke D, Esser C et al (2011). Natural aryl hydrocarbon receptor ligands control organogenesis of intestinal lymphoid follicles. *Science* (New York, NY) 334: 1561–1565.

Li Z, Zhang W, Su L, Huang Z, Zhang W, Ma L et al (2022). Difference analysis of intestinal microbiota and metabolites in piglets of different breeds exposed to porcine epidemic diarrhea virus infection. Frontiers in microbiology 13: 990642.

Lindemans C, Calafiore M, Mertelsmann A, O’Connor M, Dudakov J, Jenq R et al (2015). Interleukin-22 promotes intestinal-stem-cell-mediated epithelial regeneration. Nature 528: 560–564.

Liu L, Narrowe A, Firrman J, Mahalak K, Bobokalonov J, Lemons J et al (2023). Lacticaseibacillus rhamnosus Strain GG (LGG) Regulate Gut Microbial Metabolites, an In Vitro Study Using Three Mature Human Gut Microbial Cultures in a Simulator of Human Intestinal Microbial Ecosystem (SHIME). *Foods* (Basel, Switzerland) 12.

Lynch S, Pedersen O (2016). The Human Intestinal Microbiome in Health and Disease. The New England journal of medicine 375: 2369–2379.

Makowski L, Chaib M, Rathmell J (2020). Immunometabolism: From basic mechanisms to translation. Immunological reviews 295: 5–14.

Mizoguchi A (2012). Healing of intestinal inflammation by IL-22. Inflammatory bowel diseases 18: 1777–1784.

Pernomian L, Duarte-Silva M, de Barros Cardoso C (2020). The Aryl Hydrocarbon Receptor (AHR) as a Potential Target for the Control of Intestinal Inflammation: Insights from an Immune and Bacteria Sensor Receptor. Clinical reviews in allergy & immunology 59: 382–390.

Pitocco D, Di Leo M, Tartaglione L, De Leva F, Petruzziello C, Saviano A et al (2020). The role of gut microbiota in mediating obesity and diabetes mellitus. European review for medical and pharmacological sciences 24: 1548–1562.

Qian M, Liu J, Zhao D, Cai P, Pan C, Jia W et al (2022). Aryl Hydrocarbon Receptor Deficiency in Intestinal Epithelial Cells Aggravates Alcohol-Related Liver Disease. Cellular and molecular gastroenterology and hepatology 13: 233–256.

Ronan V, Yeasin R, Claud E (2021). Childhood Development and the Microbiome-The Intestinal Microbiota in Maintenance of Health and Development of Disease During Childhood Development. Gastroenterology 160: 495–506.

Rooks M, Garrett W (2016). Gut microbiota, metabolites and host immunity. Nature reviews Immunology 16: 341–352.

Satoh-Takayama N, Vosshenrich C, Lesjean-Pottier S, Sawa S, Lochner M, Rattis F et al (2008). Microbial flora drives interleukin 22 production in intestinal NKp46+ cells that provide innate mucosal immune defense. Immunity 29: 958–970.

Su X, Gao Y, Yang R (2022). Gut Microbiota-Derived Tryptophan Metabolites Maintain Gut and Systemic Homeostasis. Cells 11.

Wang J, Gao M, Cheng M, Luo J, Lu M, Xing X et al (2024). Single-Cell Transcriptional Analysis of Lamina Propria Lymphocytes in the Jejunum Reveals Innate Lymphoid Cell-like Cells in Pigs. Journal of immunology (Baltimore, Md: 1950).

Xia P, Liu J, Wang S, Ye B, Du Y, Xiong Z et al (2017). WASH maintains NKp46 ILC3 cells by promoting AHR expression. Nature communications 8: 15685.

Xue M, Zhao J, Ying L, Fu F, Li L, Ma Y et al (2017). IL-22 suppresses the infection of porcine enteric coronaviruses and rotavirus by activating STAT3 signal pathway. Antiviral research 142: 68–75.

Yan F, Liu L, Cao H, Moore D, Washington M, Wang B et al (2017). Neonatal colonization of mice with LGG promotes intestinal development and decreases susceptibility to colitis in adulthood. Mucosal immunology 10: 117–127.

Yang S, Li Y, Wang B, Yang N, Huang X, Chen Q et al (2020a). Acute porcine epidemic diarrhea virus infection reshapes the intestinal microbiota. Virology 548: 200–212.

Yang S, Li S, Lu Y, Jansen C, Savelkoul H, Liu G (2023). Oral administration of Lactic acid bacteria inhibits PEDV infection in young piglets. Virology 579: 1–8.

Yang W, Yu T, Huang X, Bilotta A, Xu L, Lu Y et al (2020b). Intestinal microbiota-derived short-chain fatty acids regulation of immune cell IL-22 production and gut immunity. Nature communications 11: 4457.

Yu L, Agirman G, Hsiao E (2022). The Gut Microbiome as a Regulator of the Neuroimmune Landscape. Annual review of immunology 40: 143–167.

Zegarra-Ruiz D, Kim D, Norwood K, Kim M, Wu W, Saldana-Morales F et al (2021). Thymic development of gut-microbiota-specific T cells. Nature 594: 413–417.

Zelante T, Iannitti R, Cunha C, De Luca A, Giovannini G, Pieraccini G et al (2013). Tryptophan catabolites from microbiota engage aryl hydrocarbon receptor and balance mucosal reactivity via interleukin-22. Immunity 39: 372–385.

Zenewicz L, Yancopoulos G, Valenzuela D, Murphy A, Stevens S, Flavell R (2008). Innate and adaptive interleukin-22 protects mice from inflammatory bowel disease. Immunity 29: 947–957.

