## Supplementary table for "AhR ligands from LGG metabolites promote piglet intestinal ILC3 activation and IL-22 secretion to inhibit PEDV infection": Supplementary Data 1.docx

**1. PEDV correlation detection plasmid and primer synthesis**

PEDV standard positive plasmid，M gene of PEDV（PEDV LJX strain），in pET-28a(+) ZB04293. Highlighted as protective bases and restriction sites.

GTCGACAAATGTCTAACGGTTCTATTCCCGTTGATGAGGTGATTCAACACCTTAGAAACTGGAATTTCACATGGAATATCATACTGACGATACTACTTGTAGTGCTTCAGTATGGCCATTACAAGTACTCTGCGTTCTTGTATGGTGTCAAGATGGCTATTCTATGGATACTTTGGCCTCTTGTGTTAGCACTGTCACTTTTTGATGCATGGGCTAGCTTTCAGGTCAATTGGGTCTTTTTTGCTTTCAGCATCCTTATGGCTTGCATCACTCTTATGCTGTGGATAATGTACTTTGTCAATAGCATTCGGTTGTGGCGCAGGACACATTCTTGGTGGTCTTTCAATCCTGAAACAGACGCGCTTCTCACTACTTCTGTGATGGGCCGACAGGTCTGCATTCCAGTGCTTGGAGCACCAACTGGTGTAACGCTAACACTCCTTAGTGGTACATTGCTTGTAGAGGGCTATAAGGTTGCTACTGGCGTACAGGTAAGTCAATTACCTAATTTCGTCACAGTCGCCAAGGCCACTACAACAATTGTCTACGGACGTGTTGGTCGTTCAGTCAATGCTTCATCTGGCACTGGTTGGGCTTTTTATGTCCGGTCCAAACACGGCGACTACTCAGCTGTGAGTAATCCGAGTTCGGTTCTCACAGATAGTGAGCTTAACTCGAG

| PEDV forward primer | GATACTTTGGCCTCTTGTGT |
| --- | --- |
| PEDV reverse primer | CACAACCGAATGCTATTGACG |
| PEDV Taqman probe | TTCAGCATCCTTATGGCTTGCATC |

**2. q-PCR primers**

| gene | forword | reverse |
| --- | --- | --- |
| IL-22(XM_021091967.1) | TCCTTCTCCTCGCCCTGTGG | AAGGTGCGGTTGGTGATGTAGG |
| IL1β(NM_001302388.2) | GTGTCTGTGATTGTGGCAAAGGAG | AGGACGATGGGCTCTTCTTCAAAG |
| TNP-α(NM_214022.1) | CCACCACGCTCTTCTGCCTAC | TTGAGACGATGATCTGAGTCCTTGG |
| LYZ1(NM_214392.2) | AGCCGCTACTGGTGTAATGATGG | AACGCCTAGTGGATCTCTGACAAC |
| ASCL2(NM_001122991.1) | GCGTGAAGCTGGTGAACTTGG | GCGTCTCCACCTTGCTCAGC |
| OLFM4(XM_003482903.4) | CTTGGCTGTGGATGAGAATGGATTG | CCGCAAGTGTGGTGTCATTGAG |
| LGR5(NM_001315762.1) | ATGACCATCGCCTACACCAAGC | AGGAGAAGGACAAGAAAGCCACTG |
| STAT3(NM_001044580.1) | GGAGAAGGACATCAGCGGTAAGAC | GTAGACCAGCGGAGACACAAGG |
| REG3a(NM_002580.3) | TCCCTGGTGAAGAGCATTGGTAAC | TCATCACATCACTGCTACTCCACTC |
| Reg3g(NM_011260.2) | ACAGAGGTGGATGGGAGTGGAG | ATTGCCTGAGGAAGAGGAAGGATTC |
| Reg3b(NM_011036.1) | GGAGGTGGATGGGAATGGAGTAAC | AGAAAGCACGGTCTAAGGCAGTAG |
| IFN-γ(NM_213948.1) | TGGTAGCTCTGGGAAACTGAATGAC | TTGATGAGTTCACTGATGGCTTTGC |
| IFN-α1(NM_214393.1) | CCAACCTCAGCCTTCCTCACG | TCATTTGTGCCAGGAGCCTCAG |
| NIFK(XM_021074506.1) | GCCGCTCTTGTCCCTGAATCC | TTGTTTCTCTGGTTGCTTGGTTGC |
| VILL(XM_003358368.5) | CGGCTCCTTCATCCAACACCTC | GCCTCCCTTCCTGTAGATGATTCC |
| EPCAM(NM_214419.1) | ATGATCCTGACTGTGACGAGAATGG | GTAGGTCCTCACTCGCTCCAAAC |
| IL17F(XM_013978026.2) | TCCCTGCTGCTGTTGATGTTGG | TGATTCTGACTGAGGATGCGGATG |
| IL17A(NM_001005729.1) | TGTCACTGCTGCTTCTGCTGAG | GTGCTCCGGTTCAAGATGTTCAAG |
| TNF(NM_214022.1) | CCACCACGCTCTTCTGCCTAC | TTGAGACGATGATCTGAGTCCTTGG |
| TIMP1(NM_213857.1) | TGCTGCTGGCTGTGAGGAATG | TGCTGCTGGCTGTGAGGAATG |
| CXCL8(NM_213867.1) | AAATACGCATTCCACACCTTTCCAC | GGCAGACCTCTTTTCCATTGACAAG |
| CXCL2(NM_001001861.2) | CTGCTGCTCCTGCTTCTAGTGG | ACCTTCAGGTCCTGGATGTTCTTG |
