## Supplementary figures and images for "AhR ligands from LGG metabolites promote piglet intestinal ILC3 activation and IL-22 secretion to inhibit PEDV infection"

### Fig. S1.jpg

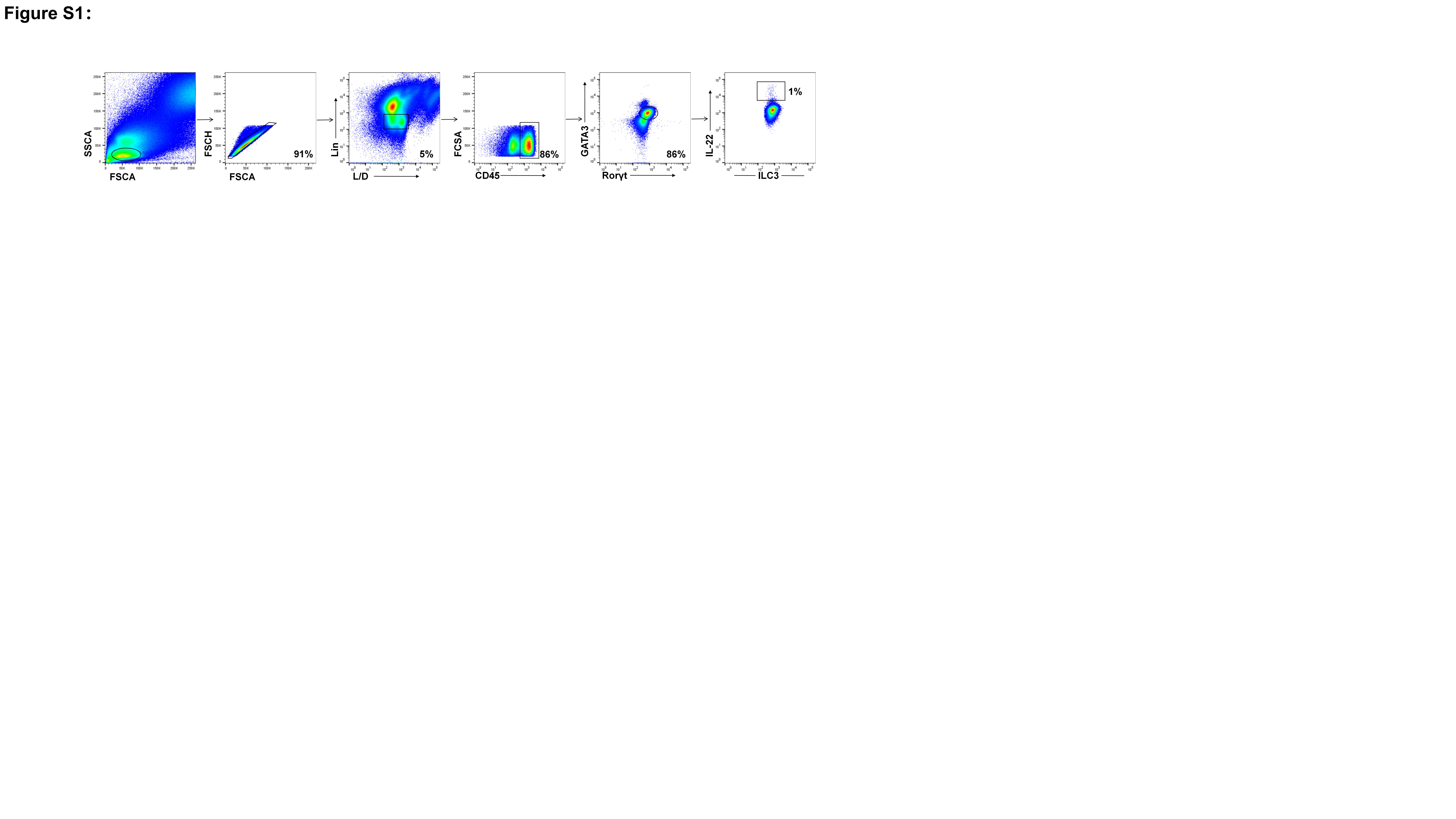

### Fig. S2.jpg

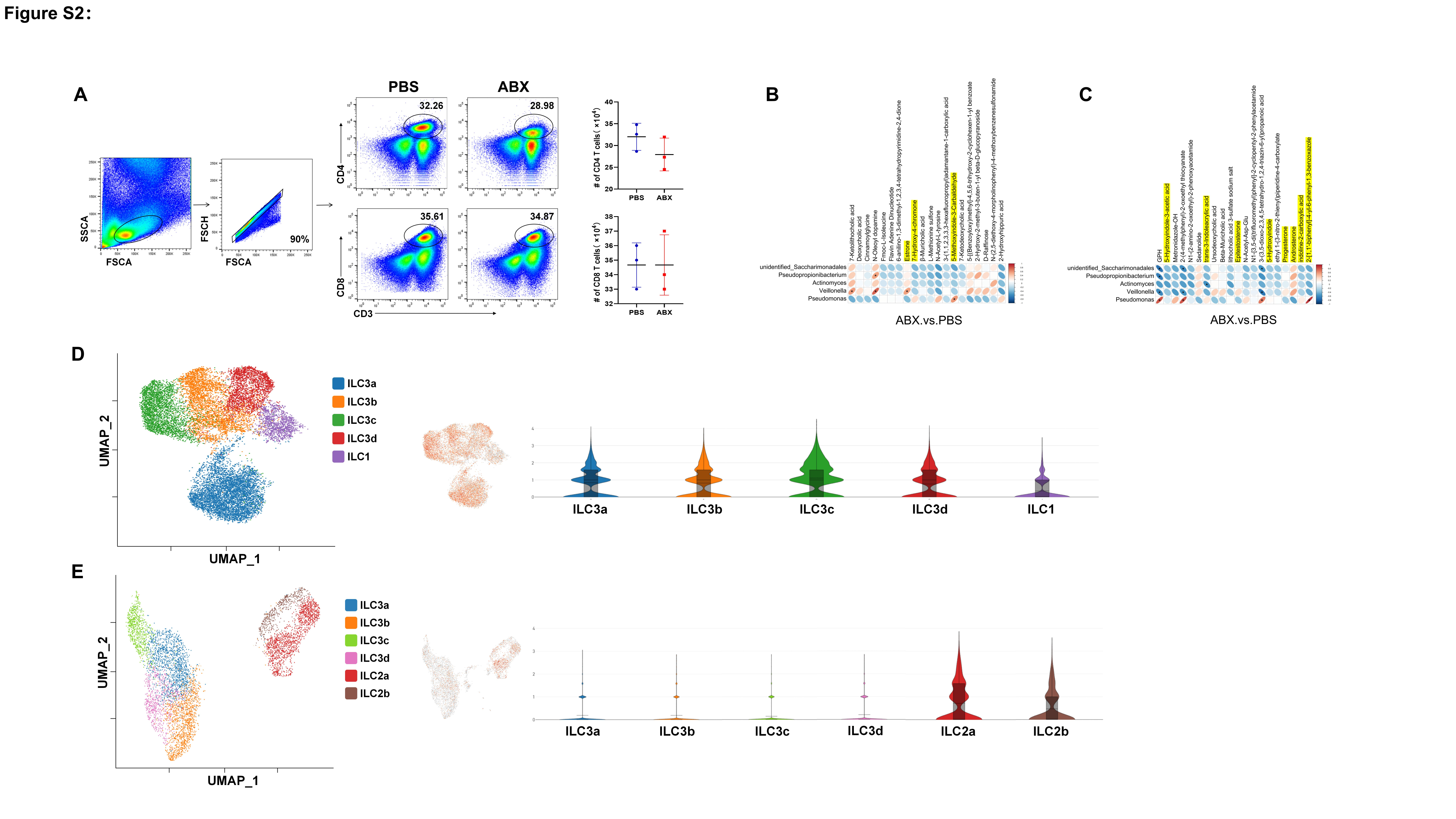

### Fig. S3.jpg

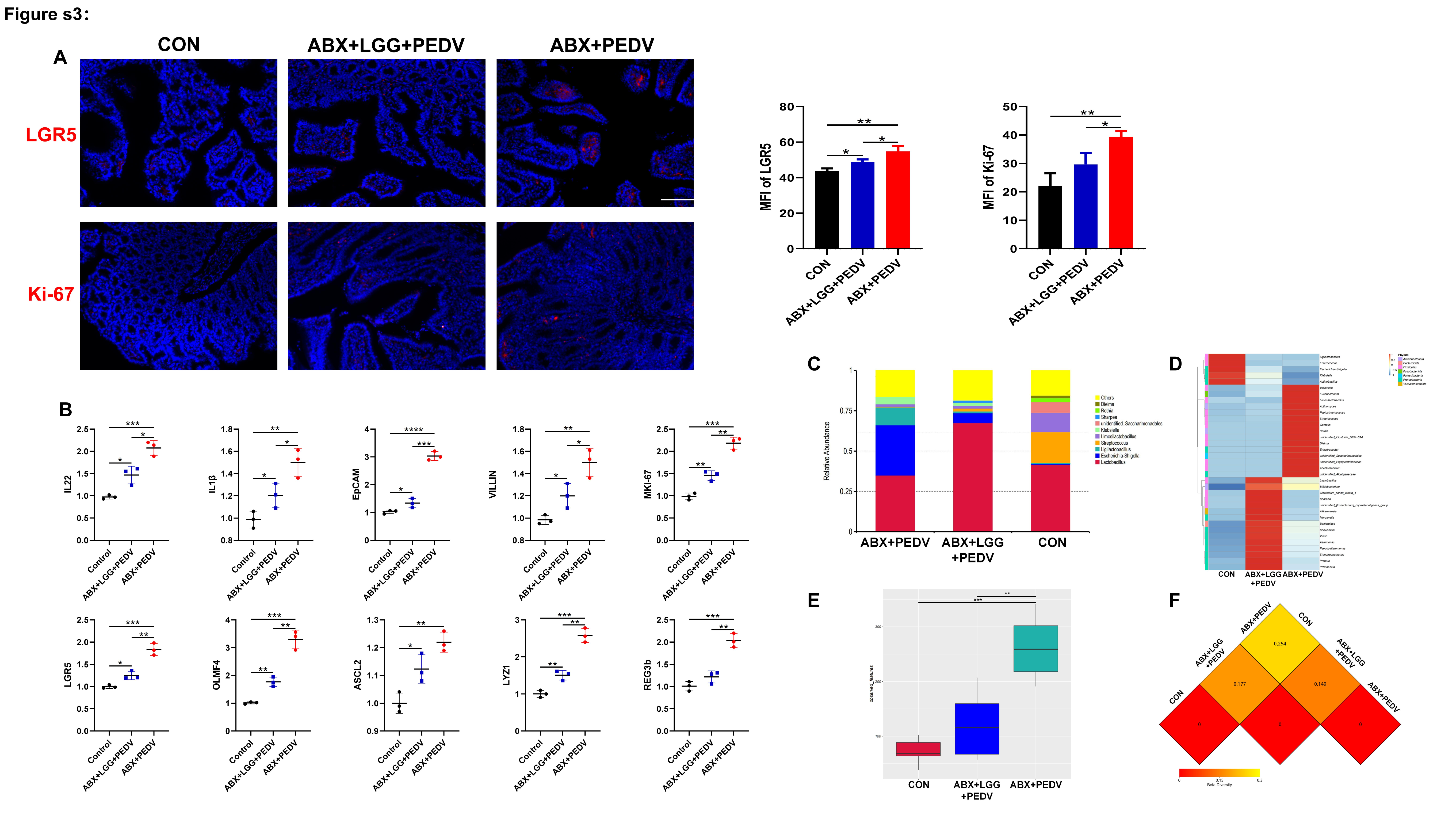

### Fig. S4.jpg

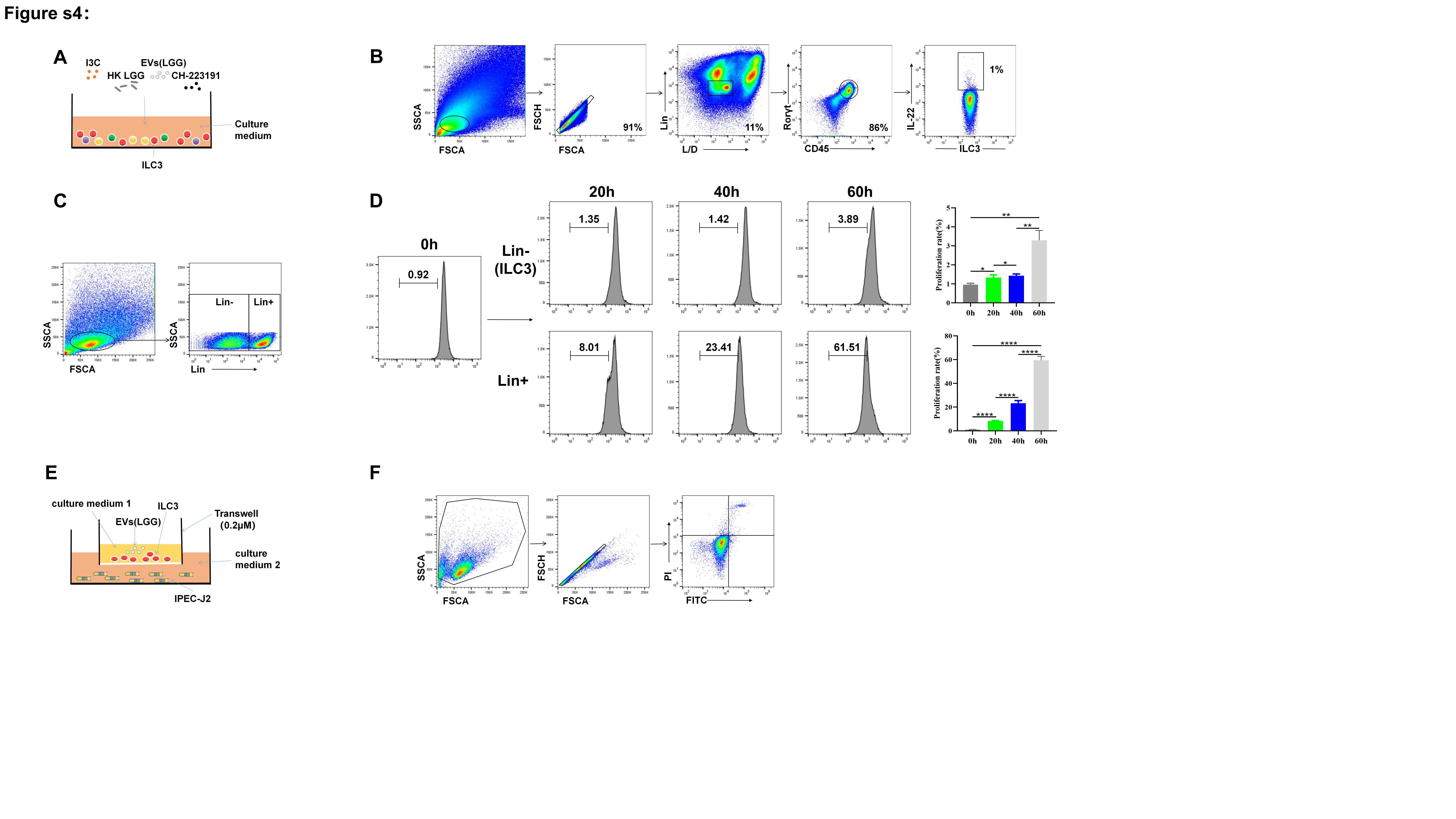

### Fig. S5.jpg

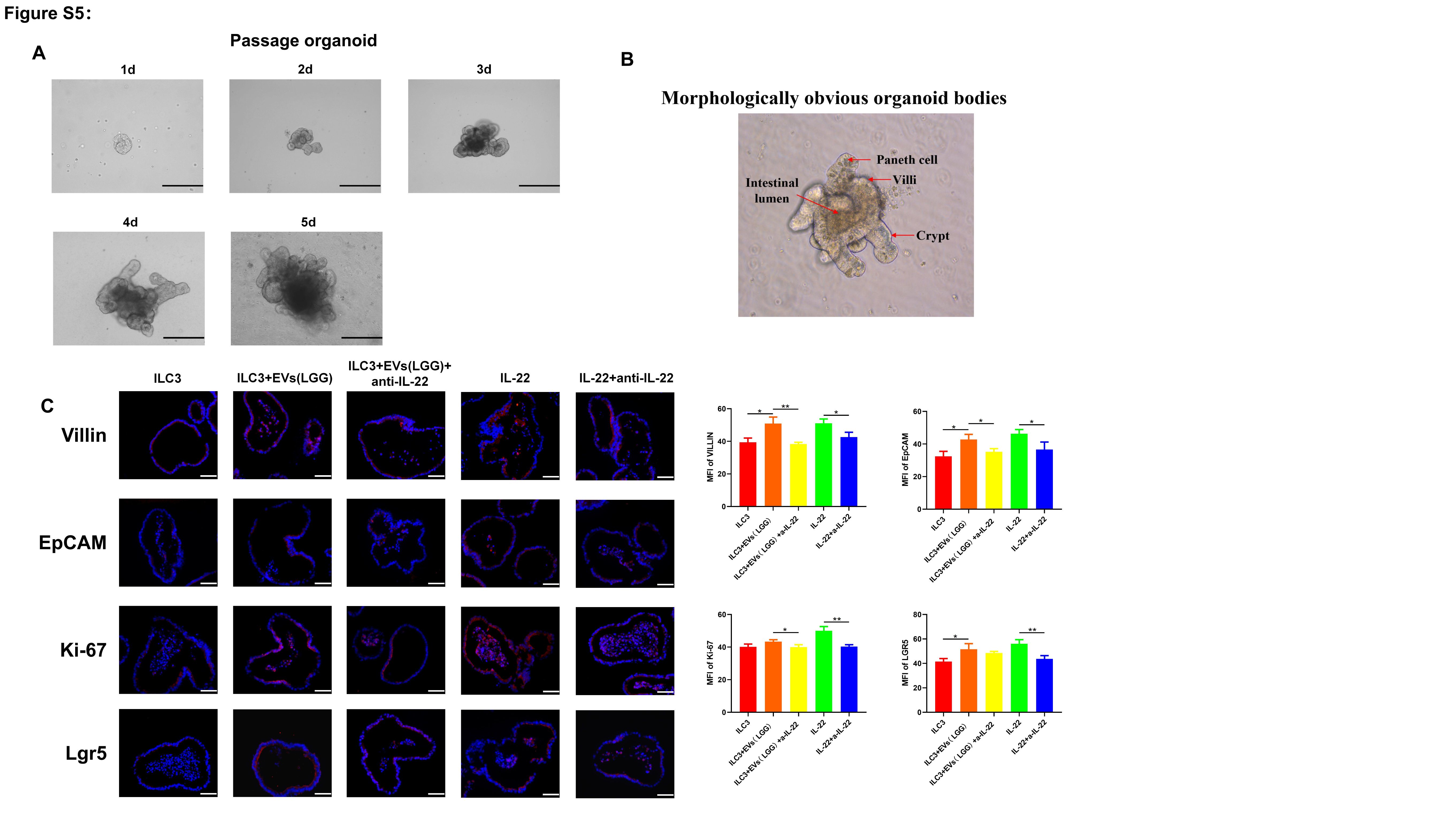
